# Exploring the druggable space around the Fanconi anemia pathway using machine learning and mechanistic models

**DOI:** 10.1101/647735

**Authors:** Marina Esteban, María Peña-Chilet, Carlos Loucera, Joaquín Dopazo

## Abstract

**Background:** In spite of the abundance of genomic data, predictive models that describe phenotypes as a function of gene expression or mutations are difficult to obtain because they are affected by the curse of dimensionality, given the disbalance between samples and candidate genes. And this is especially dramatic in scenarios in which the availability of samples is difficult, such as the case of rare diseases.

**Results:** The application of multi-output regression machine learning methodologies to predict the potential effect of external proteins over the signaling circuits that trigger Fanconi anemia related cell functionalities, inferred with a mechanistic model, allowed us to detect over 20 potential therapeutic targets.

**Conclusions:** The use of artificial intelligence methods for the prediction of potentially causal relationships between proteins of interest and cell activities related with disease-related phenotypes opens promising avenues for the systematic search of new targets in rare diseases.

## Background

With the extraordinarily fast increase in throughput that sequencing technologies underwent in the last years [1, 2], genomics has become a *de facto* Big Data discipline. Recent prospective studies have compared genomic data generation with other major data generators such as astronomy, twitter and youtube and have concluded that genomics is either on par with or, possibly even most demanding than the Big Data domains analyzed in terms of data acquisition, storage, distribution, and analysis of data [3]. Therefore, this seems to be the ideal scenario for the application of machine learning techniques, that have recently been successfully applied to many domains of medicine [4] such as radiology [5], pathology [6], ophthalmology [7], cardiology [8], etc. However, in the case of human genomic data, most of the applications have been unsupervised class discovery approaches, using gene expression data for visualization, clustering, and other tasks, mainly in single-cell [9, 10] or cancer [11, 12], being supervised applications restricted to a few examples of relatively simple problems, in which a good balance between variables to predict and data available is satisfactory, such as inferring the expression of genes based on a representative subset of them [13] or predicting the activity status of Ras pathway in cancer [14]. Consequently, in spite of the wealth of genomic data available there is a lack of translational applications due to the fact that the most interesting predictive scenarios face a serious problem of potential overfitting. Thus, attempts to describe complex, multivariant phenotypes as a function of an undefined number of genes are hampered by the high number of variables (in the range of 20,000 genes [15]), which challenge many conventional ML approaches. Therefore, new strategies that exploit the enormous potential of ML applied to genomic Big Data in order to model diseases and discover new therapies are necessary.

An especially interesting use of genomic data is related with the application of ML to model the function of the cell [16]. Such models form a natural bridge from variations in genotype (at the scale of gene activities) to variations in phenotype (at the scale of cells and organisms) [17, 18]. Despite, these models are based on yeast, an organism far simpler than human, and use yeast genomic data, which are far more abundant than human genomic data, the framework proposed is interesting not only because of the use of a causal link between genotype and phenotype but also because it is attained with a dimensionality reduction. Thus, mechanistic models of human cell signaling [19] or cell metabolism [20] can provide the functional link between the gene-level data available (gene expression) and the cell phenotype level, allowing the selection of specific disease-related cellular mechanisms of interest. In fact, mechanistic models have helped to understand the disease mechanisms behind different cancers [21–24], the mechanisms of action of drugs [19], and other biologically interesting scenarios such as the molecular mechanisms that explain how stress-induced activation of brown adipose tissue prevents obesity [25] or the molecular mechanisms of death and the post-mortem ischemia of a tissue [26].

Here we plan to use a mechanistic model of the molecular mechanism of a disease, Fanconi anemia (ORPHA:84), a rare condition that causes genomic instability and a range of clinical features that include developmental abnormalities in major organ systems, early-onset bone marrow failure, and a high predisposition to cancer [27]. Signaling is known to play a relevant role in the disease and also defines its most characteristic hallmark: failure of DNA repair [28, 29]. In addition, it has been described that FA influences survival and self-replication of hematopoietic cells [30]. Currently a detailed map of FA signaling is available in KEGG (03460) that can be used to derive a mechanistic model that relate gene expression to the activity of signaling circuits within the FA pathway that trigger cell activities related to FA hallmarks. These models can be used to investigate other molecules that could affect the activity of such circuits and therefore, presumably, to FA hallmarks. Therefore, these molecules are potential therapeutic targets. Since we are dealing with a rare disease, which typically are not considered as attractive business niches by pharmaceutical companies [31], we will restrict the search space to proteins that are already targets of approved drugs. Actually, here we are aiming for drug repurposing, that is, the discovery of new indications for drugs already used in the treatment of other diseases [32], an ideal strategy for rare diseases that accelerates enormously the evaluation of candidate molecules and simultaneously reduces failure risks. The attainment of the relationships between candidate proteins for a new indication and the FA hallmarks poses a challenge that can be addressed with the appropriate ML method.

## Results

### General approach

Here we take advantage of the biological knowledge available on FA, as represented in the FA pathway. The FA pathway describes the functional interaction among genes that finally trigger, from six different circuits, cell functionalities related with DNA repair (see Figure 1), a known FA hallmark. Since the disease condition involves the malfunction of one or several of these DNA repair cell functionalities, we hypothesize here that other genes that have an influence on the status of these functionalities might be playing the role of upstream regulators and therefore their potential modulator capacity could eventually make of them suitable therapeutic targets. In order to find druggable genes that could be playing a significant modulator role over FA hallmarks we use known drug target (KDT) genes listed in DrugBank [33] (Additional File 1).

**Figure 1.**
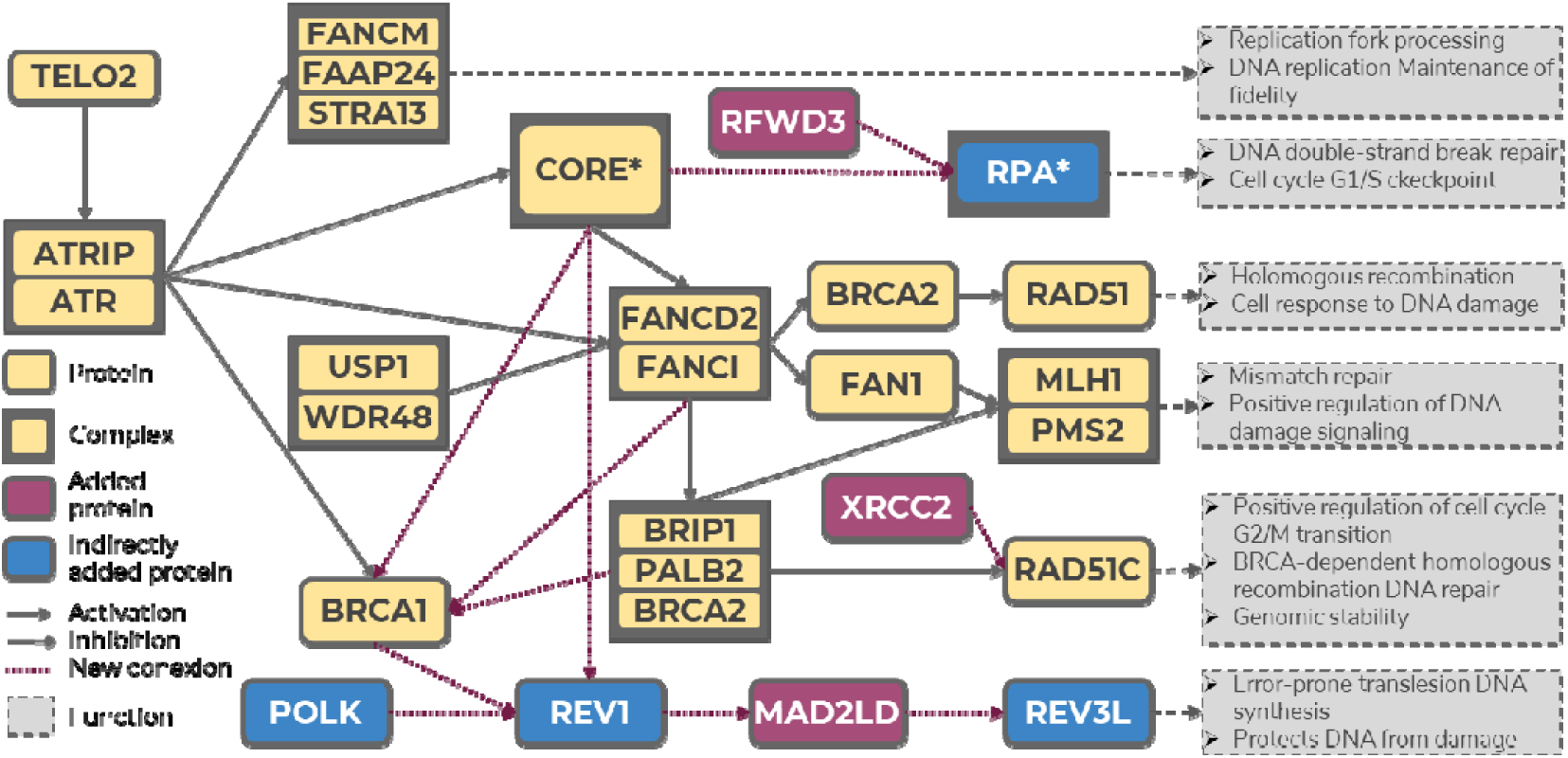
Fanconi anemia curated map, based in the KEGG FA pathway. There are two protein complexes: RPA, composed of *RPA1, RPA2, RPA3* and *RPA4*, and Core, composed of *FANCM, FANCG, FANCL, FAAP100, FANCA, FANCB, UBE2T, STRA13, FANCC, FAAP24, HES1, FANCE, FANCF, BLM, RMI1, RMI2* and *TOP3A*. At the end of the effector nodes, whose names are taken for the circuits, a description of the main functionalities triggered by the signaling circuits can be found.

These genes are used to predict the activity of the signaling circuits triggering the FA hallmarks. Since the FA pathway available in KEGG seems incomplete we first build a curated expanded version of the FA pathway (see below). Then, we search for potential known drug targets that affect the functionality of the FA pathway. Figure 2 summarizes the procedure followed: for each sample of each tissue available for each individual (over 11,000), the activity of the genes in the pathway is used to estimate the activity of the circuits contained in the FA pathway using *Hipathia* [21]. Then, across the 11,000 samples, the ML procedure tries to infer the circuit activities from the expression levels of the KDT genes external to the pathway.

**Figure 2.**
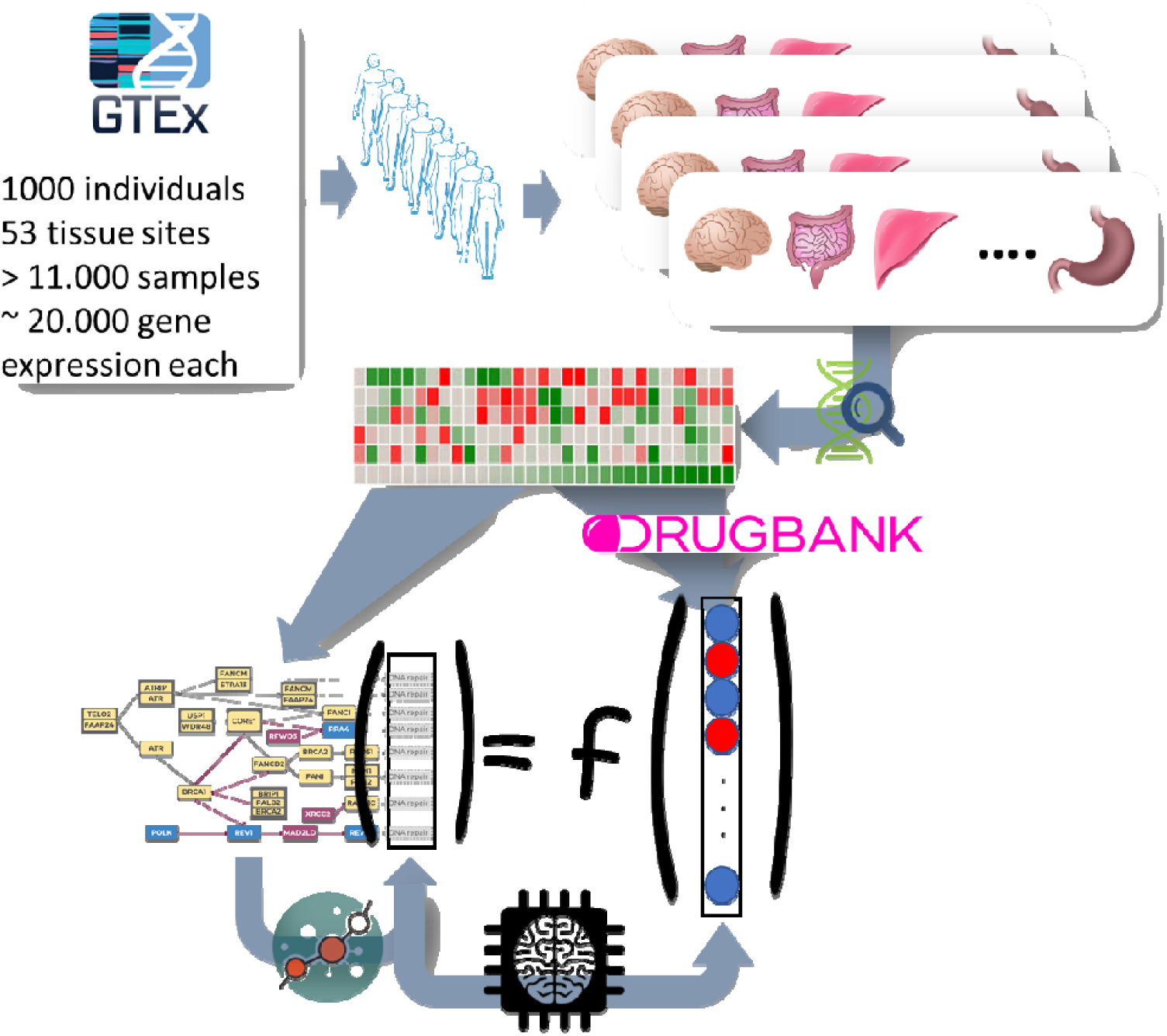
Schema of the procedure followed for the analysis.

### Building a curated Fanconi Anemia disease map

Here we use as starting point the KEGG FA pathway (hsa03460). However, among the 54 genes present in the pathway (see Additional File 2), three known FA genes (*MAD2L2, RFWD3* and *XRCC2*) described in Orphanet (ORPHA:84) were missing, which suggests that the FA KEGG pathway probably does not constitute an updated version of the current knowledge on FA. Therefore, we have derived a manually curated expanded version of the FA map. To achieve so we have used the package *pubmed.mineR* [34] with all the possible pairs of FA genes searching for direct functional interactions. The results confirmed all the gene-gene interactions described in the KEGG pathway and expanded the connections to the three genes not present in the KEGG version as well as discovered 12 new interactions among FA genes (see Table 1). Figure 1 depicts the FA pathway expanded by manual curation.

**Table 1.** New genes and connections discovered that allow the expansion of the FA pathway. The first two columns correspond to the two interactor proteins, the third column refers to the type of interaction and the last column shows the supporting bibliographic evidence. Genes *MAD2L2, RFWD3* and *XRCC2* (in bold) did not appear in the original FA KEGG pathway and were added to the new curated FA pathway

| NODE 1 | NODE 2 | INTERACTION | REF. |
| --- | --- | --- | --- |
| **MAD2L2** | *REV3L* | binding | [35] |
| **RFWD3** | *RPA1* | binding/association | [36] |
| **XRCC2** | *RAD51C* | activation | [37] |
| *REV1* | **MAD2L2** | binding/association | [35] |
| *FANCC* | *REV1* | activation | [38] |
| *POLK* | *REV1* | binding/association | [39] |
| *BRCA1* | *REV1* | activation | [40] |
| *BRIP1* | *BRCA1* | binding/association | [41] |
| *PALB2* | *BRCA2* | binding/association | [42] |
| *PALB2* | *BRCA1* | binding/association | [43] |
| *FANCA* | *BRCA1* | binding/association | [44] |
| *FANCD2* | *BRCA1* | binding/association | [45] |

Interestingly, in spite of the small number of samples in the comparison, the use of a mechanistic model, built in *Hipathia* [46] with the curated FA pathway, to analyze an experiment that compares gene expression in bone marrow cells between normal volunteers and FA patients [30] (GSE16334) rendered a significantly different activity in two circuits: REV3L (FDR-adj. p-value= 5.1×10 ^-4^) and the RPA complex (FDR-adj. p-value= 4.5×10 ^-3^), as well as the MLH1-PMS2, almost significant (see Table 2) that could not be detected when using the original KEGG FA pathway. Therefore, the curated pathway demonstrates a better detection of the expected differential behavior between normal and diseased bone marrow tissue than the original FA pathway, directly taken from KEGG. Figure 3 shows the distributions of the activities of different FA pathway signaling circuits in healthy and FA bone marrow cells in which more pronounced differences in circuit activity can be visualized for the above-mentioned circuits (REV3L, the RPA complex and MLH1-PMS2). Actually, Additional File 3 shows the same distribution obtained for the original FA KEGG pathway, where some incoherence can be observed, such as the absence of activity in four of the seven circuits. Figure 4 shows the activity in different normal tissues, taken from GTEx, which include blood, a tissue affected by the disease, two tissues with a high rate of cell replication (skin and gastrointestinal), where DNA reparation is expected to play a relevant role, and another tissue with low rate of cell replication (brain). Unfortunately, there are no expression data for bone marrow, the main tissue affected by the disease, in GTEx. DNA reparation circuits show a slightly different activity in brain when compared to the rest of tissues in the case of the three FA circuits.

**Table 2.** Differential circuit activity in a comparison of healthy versus FA bone marrow cells. Circuits are named after their effector nodes (see Figure 1)

| CIRCUIT | Activation | Statistic | p-value | FDR adj. p-value |
| --- | --- | --- | --- | --- |
| RAD51 | UP | 0.615 | 0.558 | 0.659 |
| MLH1-PMS2 | UP | 2.400 | 0.016 | 0.067 |
| REV3L | DOWN | -3.789 | $3.917 \times 10^{-5}$ | $5.092 \times 10^{-4}$ |
| RAD51C | DOWN | -1.924 | 0.056 | 0.162 |
| RPA* | UP | 3.412 | $6.923 \times 10^{-4}$ | $4.500 \times 10^{-3}$ |
| FANCM-STRA-FAAP24 | UP | 1.885 | 0.062 | 0.162 |

**Figure 3.**
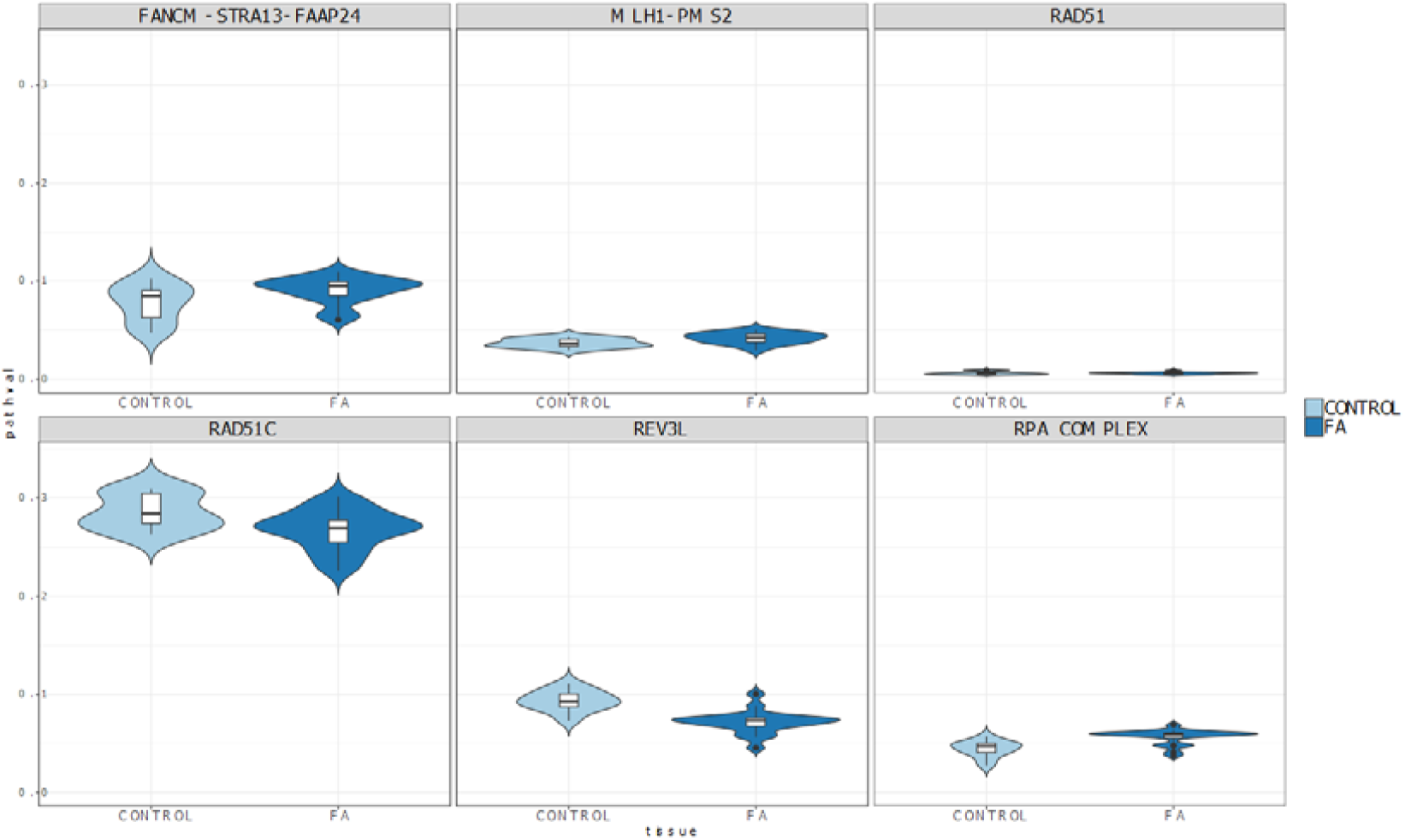
Observed distribution of circuit activities in the comparison between healthy and FA bone marrow cells.

**Figure 4.**
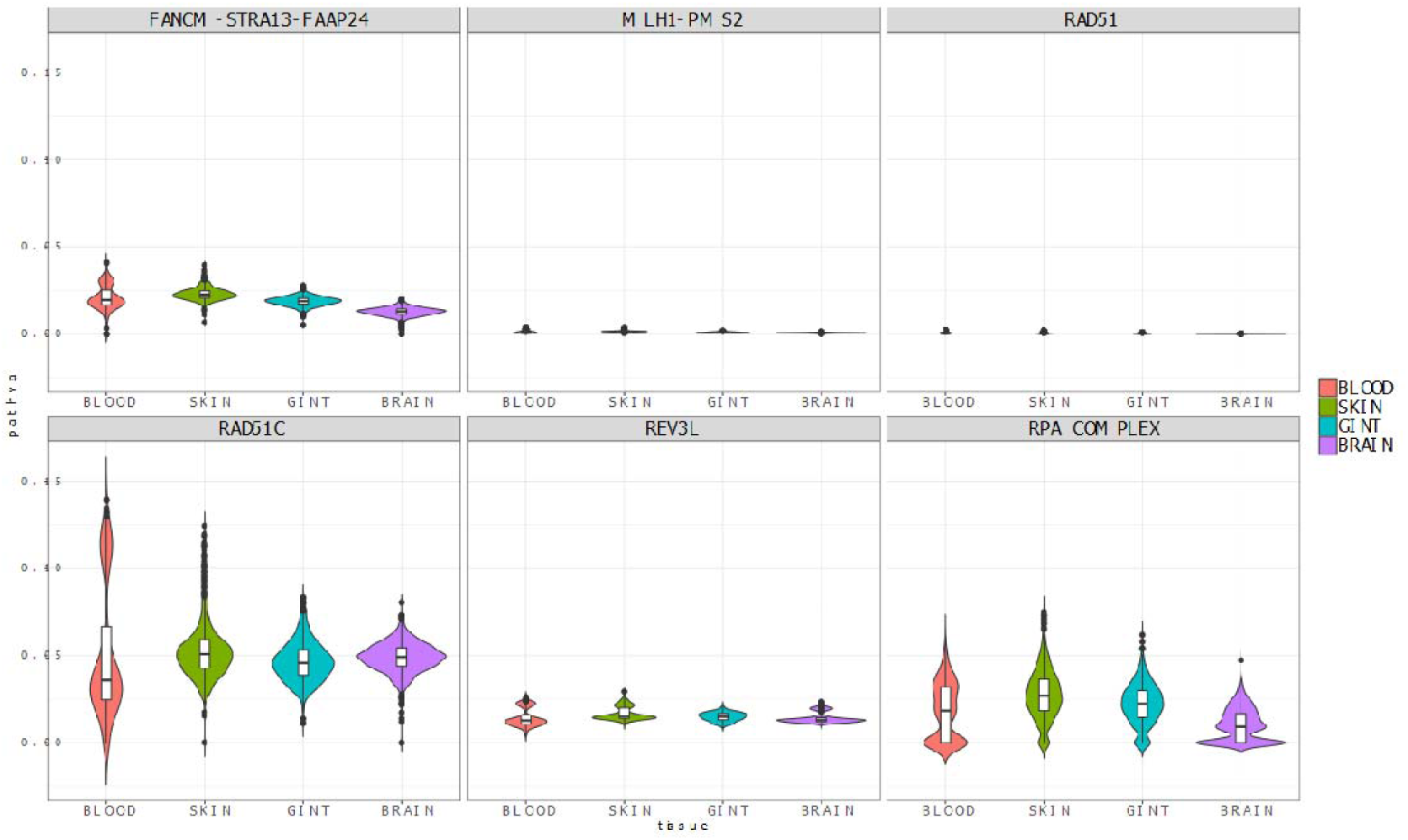
Observed distribution of circuit activities in blood, a tissue affected by the disease, two tissues with a high rate of cell replication (skin and gastrointestinal), where DNA reparation is expected to play a relevant role and another tissue with low rate of cell replication (brain).

### Exploring the druggable space of influence over the FA pathway

As sketched in Figure 2, the ML strategy was applied to detect proteins whose activity was able of predicting the activity of the FA circuits that trigger the FA hallmarks. The initial search space was restricted to KDTs extracted from DrugBank (See Additional File 1). The cross-validation of the relevance values (Figure 5) rendered a threshold of 0.006, above which the most relevant genes presented a stable value.

**Figure 5.**
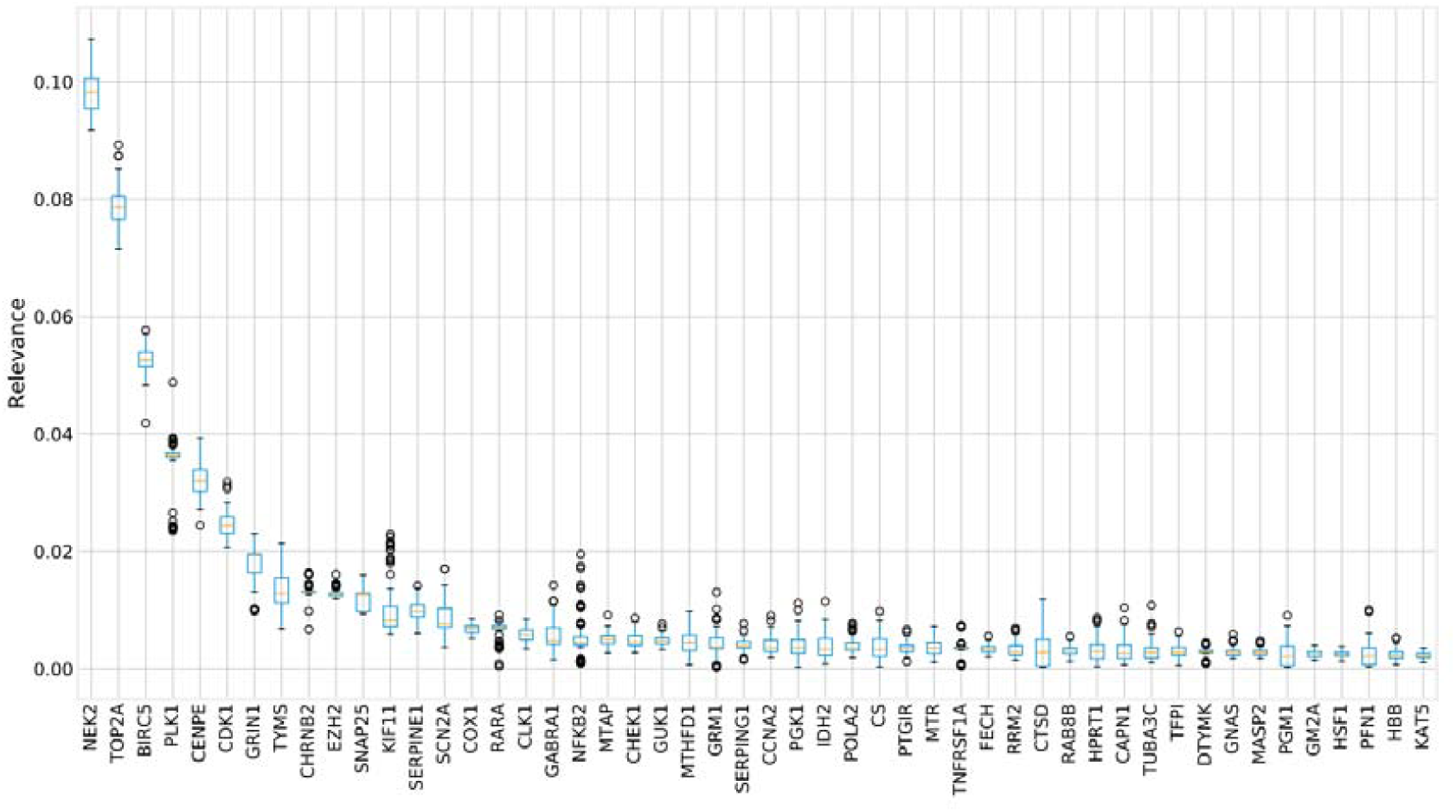
Distributions of the cross-validation of the relevance values for the top 50 most relevant genes ordered by their mean. Above the relevance value of 0.006 the relevance rendered by the ML procedure and the means obtained from the cross-validation are consistent. Then this value is taken as a threshold.

The importance of the genes selected by the ML strategy is strongly supported by a high predictive performance across all the splits, as can be seen in Figure 6. The distribution of the R^2^ score for each signaling circuit of the FA curated pathway across all the training/test splits have in all the cases a value close to 1 (note that the R^2^ score goes from -infinite to 1, where 0 represents a model that always predicts the mean for each task and a perfect model has a score of 1).

**Figure 6.**
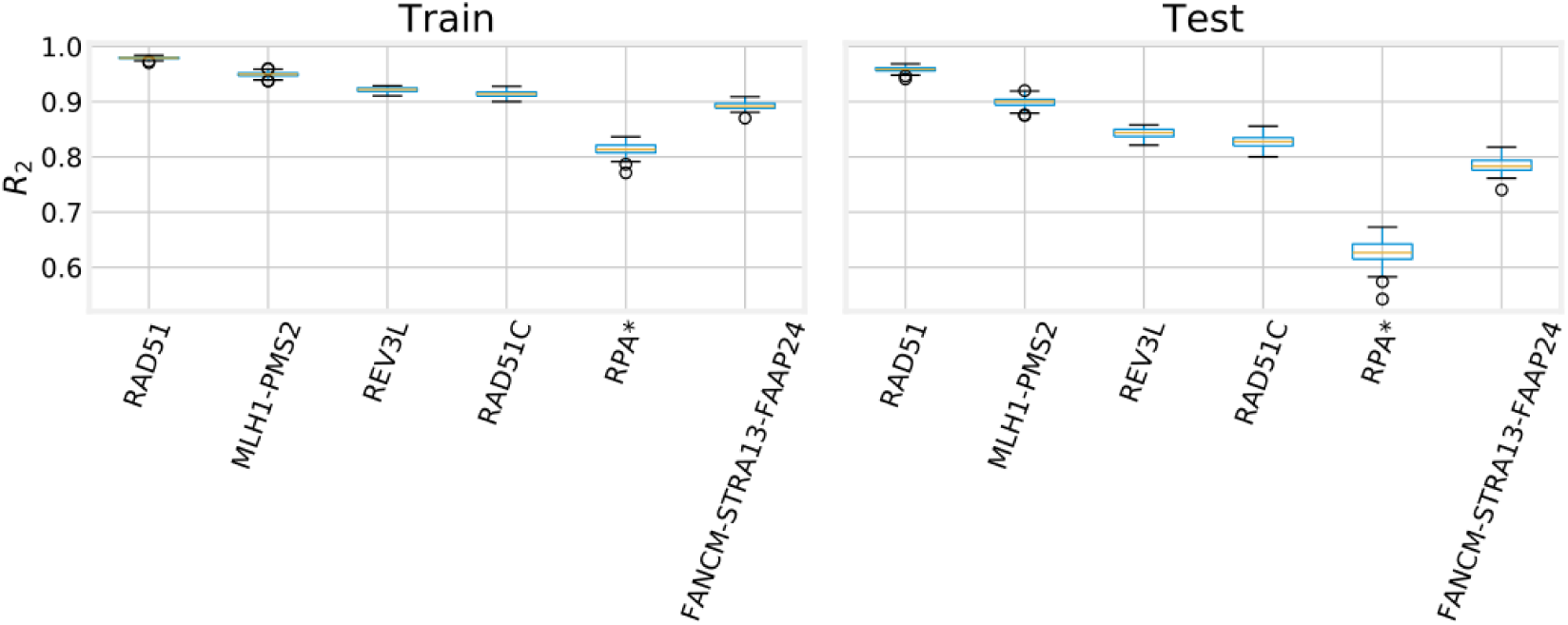
the distribution of the R_2_ score for each signaling circuit of the FA pathway across all the training/test splits. The R_2_ score goes from -infinite to 1, where 0 represents a model that always predicts the mean for each task and a perfect model has a score of 1.

A total of 17 genes resulted to have a relevance over the 0.006 threshold (See Table 3). Additional File 4 contain details on the drugs targeting these proteins.

**Table 3.**
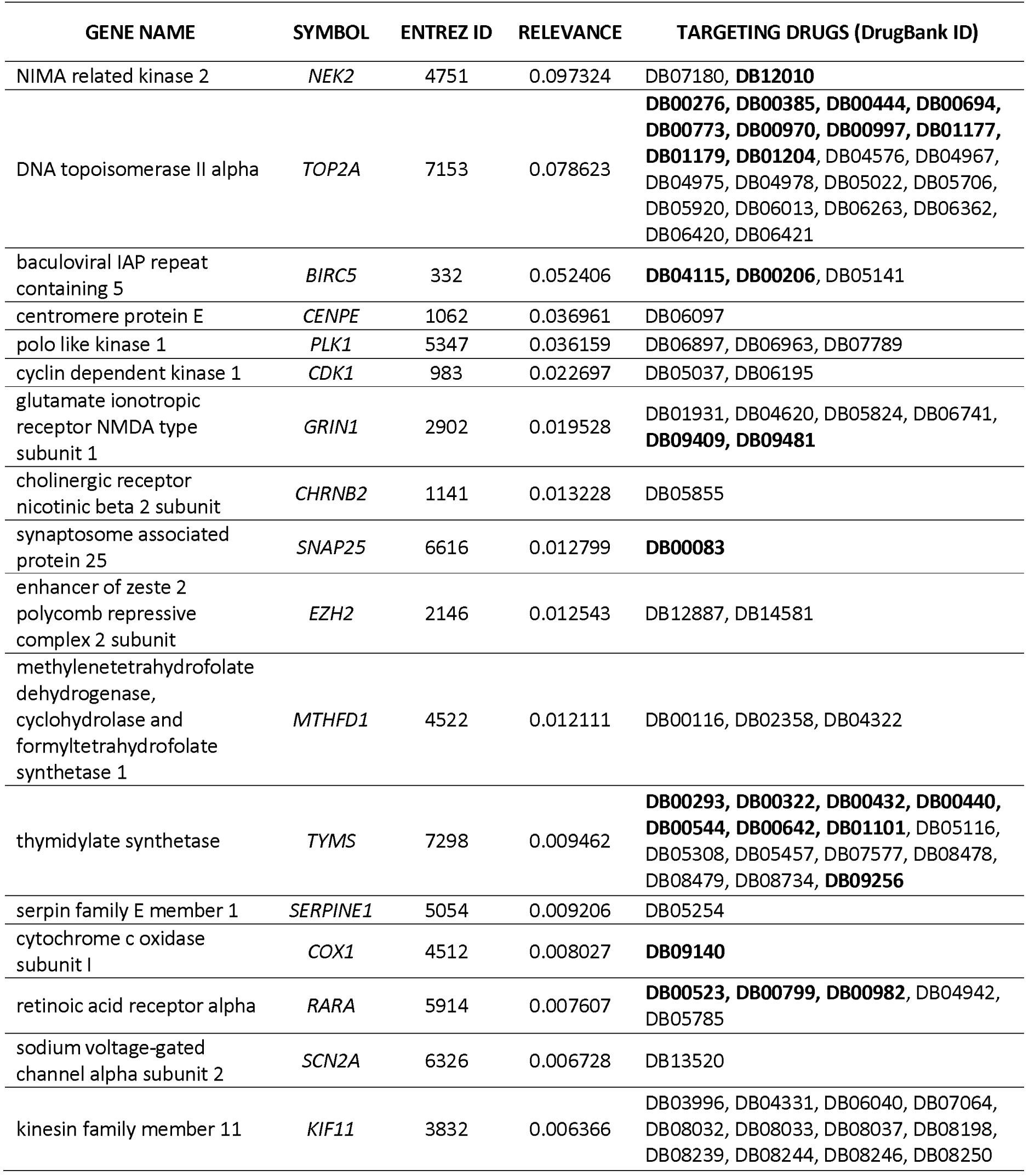
List of most relevant genes (relevance > 0.006) obtained by the model. Drug IDs in bold are approved for use according to DrugBank database

## Discussion

### Mechanistic models and Machine learning approach used

Supervised ML applications in the case of human genomic data aiming to find genes potentially causal of phenotypes have restricted to a few cases in quite simple scenarios, such as the inference of very simple (and univariate) phenotypes, such as the activity status of Ras pathway in cancer [14]. Here we aimed to approach the pathologic phenotype problem in more detail, trying to capture the complexity of the molecular mechanism of the disease. To achieve so, we have used signaling circuit activities inferred by mechanistic models, as proxies of disease-related cell functionalities triggered by them. Such mechanistic models use gene expression data to produce an estimation of profiles of signaling or metabolic circuit activity within pathways [20, 24] and have been used to describe the molecular mechanisms behind different biological scenarios such as the explanation on how stress-induced activation of brown adipose tissue prevents obesity [25], the common molecular mechanisms of three cancer-prone genodermatoses [47] or the molecular mechanisms of death and the post-mortem the ischemia of a tissue [26]. Moreover, recent benchmarking of mechanistic modeling methods shows how *Hipathia* clearly outperform to other competing method [48].

To assess the suitability of the expanded FA pathway, we have analyzed the distribution of the activity of its circuits once modeled in *Hipathia*. As expected, the overall activity in blood, skin and gastrointestinal tissues is higher than that of brain cells, due to its higher replication rate (Figure 4). However, brain tissue also exhibits pathway activity to some extent, which can be explained by the involvement of FA pathway in DNA repair, since brain cells have high level of metabolic activity and use distinct oxidative damage repair mechanisms to remove DNA damage [49]. We also observed in Figure 3 that RAD51C and REV3L circuit activities derived from the expanded FA pathway are, contrarily to the results obtained from KEGG FA pathway (Additional Figure 3), significantly lower in FA patients than in healthy donors. This observation is coherent with the fact that these circuits are involved in DNA crosslinking repair during homologous recombination, a mechanism that has been demonstrated to be damaged in FA patients [37].

Therefore, the mechanistic models of the extended FA pathway offer the possibility of discovering what protein activities potentially affect the different pathway activities that trigger FA hallmarks, which provide a mechanistic link between such proteins and the disease phenotype. However, finding these relationships constitutes a complex problem that involve multiple variables (here KDT proteins) to predict multiple outputs (here signaling circuit activities related with DNA repair, a FA hallmark) that can be formulated as multi-output regression problems (MOR), also called multi-task learning or vector valued regression. MOR is a fundamental problem in machine learning as it deals with the ability to predict multivariate responses with a single model, instead of learning one model per output, the classic single output regression (SOR) scenario, e.g. conventional univariate regression. The MOR scenario has several advantages over SOR: on the one hand, in SOR, each variable to predict is treated as independent (uncorrelated). Actually, a different set of hyper-parameters (i.e. a different model) is needed for each variable, leading to several training/testing/validation scenarios with different features learned. On the other hand, in the MOR learning framework a unique model (only one set of hyper-parameters) is used to predict all the output variables at once, with the ability to exploit and learn the shared patterns between them. Therefore, the MOR scenario provides an ideal framework to properly address hypothesis from a systems biology point of view given that it assumes that the response variables, here the different signaling circuits in the FA pathway are (or can be) interconnected. An additional advantage of using mechanistic models is that, by accurately defining the functional space of interest (the FA hallmarks described in the FA pathway), the number of circuits involved in their activity results relatively low, which constitutes a reduction of the dimensionality of the output space based on biological knowledge.

Here we used Random Forests (RF) [50], an ensemble of decision trees that aggregates the output of each estimator in order to stabilize and improve the prediction power. RFs and other tree-based ensembles have been proven to be extremely well suited for interpretable machine learning across different systems biology scenarios [51]. Tree-structured methods (TSM) provide a set of interpretable rules by splitting data into sample/target-wise homogenous groups and averaging the results. However, the predictive performance of a single decision tree is subpar when compared to other methods, such as Support Vector Machines, mostly due to the fact that a tree must make several sequential choices based on a subset of the data and one incorrect decision can impact the rest of the sequence, thus propagating the error. To improve the performance of a decision tree, several strategies have been proposed, the most notable among them are those based on building an ensemble of trees, where several trees (from hundreds to thousands) are fitted on different partitions of the training data or under different conditions, and then combined in order to achieve a better prediction capability [52]. On top of this, RFs are particularly well suited for the analysis of genomics datasets [53, 54] due to its robustness in scenarios affected by the curse of dimensionality.

Although one key advantage of RFs is its ability to produce good enough results with minimal hyperparameter search (given a sufficiently large number of trees are trained), in some circumstances the hyperparameter space must be properly optimized in order to obtain a good set of results [55]. Our problem setup is one of such cases, where a large number of highly correlated predictor variables (gene expression) interact with a multivariate response with many self-interactions (pathway circuit activities). To overcome such difficulties, we make use of Tree-structured Parzen Estimator (TPE) [56], a Sequential Model-based Global Optimization strategy for hyperparameter optimization. The base learners of a RF, the decision trees, can be easily extended to the multi-output scenario [57] by introducing a covariance weighting to the splitting criterion with the aim of finding a representation of homogeneous clusters with respect to both the predictor and response spaces. This multivariate splitting function leads to a natural extension of the relevance scores, which maintains the interpretability.

Thus, interpretability in TSM methods depends in the last instance of relevance scores, which are computed for each input variable (gene expression in our case) by averaging the importance measure (the higher, the better) of each individual tree. Recent studies [58] have concluded that, by means of the averaging of relevance technique, RF could deliver an unreliable importance measure in certain situations, such as classification problems, where the input space has many categorical variables, favoring those variables with a higher number of categories. Although here, predictor and response variables are continuous, multivariate regression is performed instead of classification, the relevance scores have been validated by studying their distribution along the repeated k-fold cross-validation methodology. Figure 5 shows the top 50 gene relevance distributions, ordered by their mean. The genes found as relevant have a significant predictive impact on the circuits as Figure 6 documents.

By means of the strategy presented here, many of the problems affecting the analysis of genomic Big Data in a ML framework can be overcome to fully exploit the discovery potential of genomic big data.

### Drugs with a potential new indication for FA

In order to understand what are the general roles played in the cell by the genes selected as most relevant by the ML algorithm (see Table 3) we carried out an enrichment analysis. The functional landscape revealed by the analysis include Gene Ontology (GO) Biological processes terms mainly related to cell cycle, specifically to the correct regulation of spindle formation, chromatin condensation, centrosome separation and in general, correct mitotic cell phase transition (see Figure 7 and Additional File 5 for a detailed description of the terms found). These terms specifically involve processes related to DNA replication, DNA repair and stress response, which suggests that the activity of these genes may potentially impact DNA repair cell ability, by controlling the balance between accumulation of mutations and apoptosis in the cell, which indirectly also impacts on tumor predisposition. Interestingly, the rare diseases most associated with relevant genes included Fanconi anemia, as well as other related diseases such as Baller-Gerold syndrome (OMIM:218600), Ataxia telangiectasia (OMIM:208900), Bloom syndrome (OMIM:210900), Filippi syndrome (OMIM:272440), Congenital aplastic anemia (OMIM:609135), Meier-Gorlin syndrome (OMIM:224690), Seckel syndrome (OMIM:606744, OMIM:210600, OMIM:613676, OMIM:613823, OMIM:614728, OMIM:615807, OMIM:616777, OMIM:617253, OMIM:614851), cutaneous melanoma (OMIM:609048). All these diseases share with FA several of its hallmarks like chromosomal instability condition or tumor predisposition [59–61].

**Figure 7.**
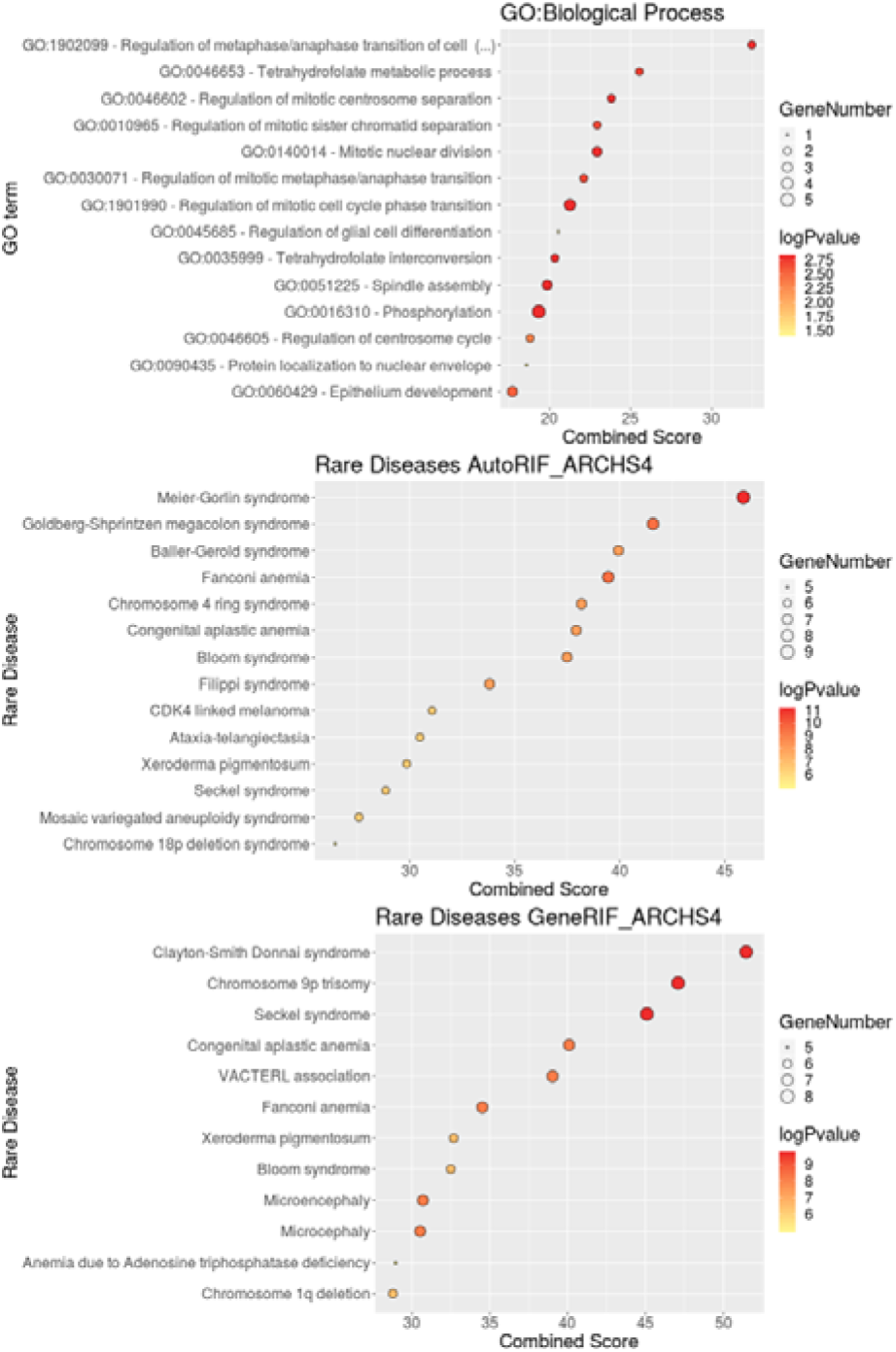
Enrichment analysis with GO terms and rare diseases.

Among the most relevant gene drug targets (Table 3) 8 proteins targeted by approved drugs, *NEK2, TOP2A, BIRC5, COX1, GRIN1, RARA, SNAP25* and *TYMS* can be found, revealing the high potential for therapeutic targets and candidates for drug repositioning in FA (Additional File 4) rendered by the ML strategy applied. Although a detailed discussion on the nature of the most relevant targets is out of the scope of this manuscript, some of the top scored ones deserve to be reviewed for their potential links to FA.

The most relevant protein, *NEK2*, is a serine/threonine-protein kinase that regulates mitosis. Its expression rises during S phase and reach its maximum level in late G2 phase, just before mitosis. The protein regulates the correct spindle formation and chromatin condensation, playing a major role in cell cycle [62]. Indeed, DNA damage results in G2 arrest due to the drastically decreasing in *NEK2* presence [63]. Indeed, this *NEK2* inhibition is dependent of ATM, a protein that, along with ATR, are master controllers of cell cycle and DNA repair, the main pathway deregulated in Fanconi Anemia [64]. *NEK2* phosphorylates FANCA, a protein conforming the FA core and highly associated with Fanconi Anemia disease [65]. These associations are in line with the expected results, supporting the robustness and suitability of the methodology presented here for the discovery of genes and new therapeutic targets relevant to diseases, FA in this case.

The protein *TOP2A* is a topoisomerase, a nuclear enzyme that binds to the DNA and alters its topologic state during transcription. It is associated with the initiation of neoplasms, such as breast and peripheral nerve tumors or Bloom syndrome, as well as with several anemia disorders (Anemia due to Adenosine triphosphatase deficiency, Congenital dyserythropoietic anemia and Congenital aplastic anemia) [66]. Regarding its connection with DNA repair, *TOP2A* show a consistent high expression in G2, but it is also highly expressed in late S phase, supporting a role in regulating entry into mitosis [67]. Besides, topoisomerase-1 and 2A gene copy numbers are elevated in patients mismatch repair-proficient tumor samples, suggesting that *TOP2A* is required to deal with high replication stress [68].

Protein *BIRC5*, also known as survivin, plays an important role in apoptosis, being involved in pathways such as *Apoptosis* (hsa04210, hsa04215), *Hippo signaling pathway* (hsa04390) and specific disease pathways such as *Pathways in cancer* (hsa05200) and *Colorectal cancer* (hsa05210). Indeed, several studies demonstrate its association with neoplasia and, specifically with colorectal cancer [69]. Some works suggests that the role of survivin in DNA repair by homologous recombination has a direct impact in cancer [70]. The gene *BIRC5* is a member of the inhibitor of apoptosis gene family (IAP), thus its downregulation promotes apoptotic cell death. One of the main mechanisms of apoptosis inhibition is due to its protection of the cell towards the action of caspases. Actually, the mechanism by which the Jak/STAT pathway specifically triggers one of the survival circuits of the apoptosis pathway that eventually results in the disease has previously been described by means of a mathematical model [71].

The protein coded by *GRIN1*, Glutamate Ionotropic Receptor NMDA Type Subunit 1, directly bind thorough NMDA receptors to their ligands (glutamate in this case) allowing calcium to enter the cell, thus, promoting cell activity and proliferation. Interestingly, some studies associate the deregulation of *GRIN1* and other NMDA receptors with tumor formation [72].

*TYMS* (Thymidylate Synthetase) protein plays a critical role in DNA replication and repair [73]. Mutations in its enhancer region, resulting in an overexpression of TYMS, are associated with several cancers and response to chemotherapy [74]. Interestingly, chemotherapeutic agents targeting *TYMS*, and reducing its expression, have grade 1 anemia as secondary effects, suggesting that deleterious mutations in this gene may produce anemia [75]. Some authors have described that *HDAC* inhibits both *TYMS* and *BIRC5* (one of the most relevant proteins found by our model), suggesting an indirect relation between both proteins [76]. But not only with *BIRC5*, a recent study showed a non-canonical interaction between *TYMS* and *FANCD2*, a protein belonging to FA pathway [77].

Gene *COX1* (Mitochondrially Encoded Cytochrome C Oxidase I) codes for the subunit 1 of Cytochrome C oxidase, the component of the respiratory chain that catalyzes the reduction of oxygen to water. Defects in this gene are associated with Acquired Idiopathic Sideroblastic Anemia (ORPHA75564), a disease that affects bone, bone marrow and myeloid tissues, phenotypes also present in Fanconi Anemia. *COX* enzymes have a role in response to oxidative stress, COX-1 is believed to play a constitutive housekeeping role [78] and its inhibition induce apoptosis and lead to Prostaglandin production induced by ionizing radiation [79]. In line with this, it has recently been demonstrated that downregulation of *COX1* stimulates mitochondrial apoptosis through NH-kB signaling pathway [80].

*RARA* (Retinoic Acid Receptor alpha) protein is involved in regulation of several cell processes, including cell differentiation, apoptosis and transcription of clock genes. Mutations in *RARA* gene, mostly resulting in fusion genes, are associated with abnormality of blood forming tissues, leukemias and deregulate genes involved in DNA repair [81]. Recent works have demonstrated in Escherichia coli that *rarA*, via its gap creation activity, generates substrates for post-replication repair pathways, including homologous recombination and translesion DNA synthesis [82], both DNA repair pathways are involved in FA disease mechanism.

With respect to the 81 drugs targeting the most relevant genes, 55 of them have a description or indication provided by DrugBank, and 28 are already approved as a therapeutic option. Of these, 37 (67.27%) drugs are indicated for cancer treatment (including breast and colorectal cancer, but mostly, leukemias), most of them have antineoplastic effects (23, 38.33%), including chemotherapeutic agents. The remaining drugs are indicated for a variety of conditions, including infections (viral or bacterial), hypertension, neuropathies, Alzheimer, schizophrenia or rheuma, acting as antinflammatory, antipsychotic, antibacterial or antiviral. Most of the obtained drugs impact in the ability of the cell to perform correct replication and division.

## Conclusions

We have demonstrated how a mechanistic model, which provide a definition of cell functionalities and outcomes that account for the phenotype of the disease, can be used in combination with ML methods and genomic big data available to discover proteins that might have influence over such disease-related cell functionalities and, most likely, on the phenotype of the disease. Depending on the specific molecular mechanism of the disease and the type of influence, the molecules found can be considered therapeutic targets.

Building an interpretable model makes possible understanding how the model learns and, consequently, a disease-centric learning framework can be built. In this way, many of the problems affecting the analysis of genomic data in a ML framework can be overcome to fully exploit the discovery potential of such Big Data.

## Methods

### Data

The FA pathway (hsa03460) was obtained from KEGG. The list of FA genes (Table 4) was taken from the Orphanet [83] database (ORPHA:84).

**Table 4.** Fanconi Anemia ORPHANET (ORPHA:84) database affected genes.

| GENE NAME | SYMBOL | ENTREZ ID | ENSEMBL ID | OMIM |
| --- | --- | --- | --- | --- |
| Fanconi Anemia complementation group F | *FANCF* | 2188 | ENSG00000183161 | 603467 |
| Fanconi Anemia complementation group C | *FANCC* | 2176 | ENSG00000158169 | 227645 |
| Breast cancer type 2 susceptibility protein | *BRCA2* | 675 | ENSG00000139618 | 114480 |
| Breast cancer type 1 susceptibility protein | *BRCA1* | 672 | ENSG00000012048 | 113705 |
| Fanconi Anemia complementation group E | *FANCE* | 2178 | ENSG00000112039 | 600901 |
| RAD51 recombinase | *RAD51* | 5888 | ENSG00000051180 | 114480 |
| Fanconi Anemia complementation group D2 | *FANCD2* | 2177 | ENSG00000144554 | 227646 |
| Fanconi Anemia complementation group M | *FANCM* | 57697 | ENSG00000187790 | 609644 |
| DNA repair protein RAD51 homolog 3 | *RAD51C* | 5889 | ENSG00000108384 | 602774 |
| Ubiquitin-conjugating enzyme E2 T | *UBE2T* | 29089 | ENSG00000077152 | 610538 |
| Fanconi Anemia complementation group B | *FANCB* | 2187 | ENSG00000181544 | 300514 |
| Fanconi Anemia complementation group G | *FANCG* | 2189 | ENSG00000221829 | 602956 |
| Fanconi Anemia complementation group I | *FANCI* | 55215 | ENSG00000140525 | 609053 |
| Fanconi Anemia complementation group L | *FANCL* | 55120 | ENSG00000115392 | 608111 |
| partner and localizer of BRCA2 | *PALB2* | 79728 | ENSG00000083093 | 114480 |
| SLX4 structure-specific endonuclease subunit | *SLX4* | 84464 | ENSG00000188827 | 613278 |
| Ring finger and WD repeat domain 3 | *RFWD3* | 55159 | ENSG00000168411 | 614151 |
| BRCA1 interacting protein C-terminal helicase 1 | *BRIP1* | 83990 | ENSG00000136492 | 114480 |
| ERCC excision repair 4, endonuclease catalytic | *ERCC4* | 2072 | ENSG00000175595 | 133520 |

| subunit |  |  |  |  |
| --- | --- | --- | --- | --- |
| Mitotic arrest deficient 2 like 2 | *MAD2L2* | 10459 | ENSG00000116670 | 604094 |
| X-ray repair cross complementing 2 | *XRCC2* | 7516 | ENSG00000196584 | 600375 |
| Fanconi Anemia complementation group A | *FANCA* | 2175 | ENSG00000187741 | 227650 |

A gene expression microarray study to identify differences at the transcription level in bone marrow cells between normal volunteers and FA patients [30] was downloaded from GEO (GSE16334) and used to check the performance of the expanded FA disease map model in a real scenario.

Gene expression data from 53 non-diseased tissue sites across nearly 1000 individuals, more than 11.000 samples and 20.000 gene expression measurements each, were downloaded from the GTEx Portal [84] (GTEx Analysis V7; dbGaP Accession phs000424.v7.p2).

Genes that are target of approved drugs were taken from the DrugBank [33] database (Version 5.1.2). A total of 965 known drug target (KDT) genes targeted by a total of 7122 drugs were considered in this study (see Additional File 1). Some of these genes may potentially affect the whole FA pathway or some of their circuits, affecting in consequence, to the cell functionalities triggered by the affected circuits.

### RNA-seq data processing

After constructing the gene expression matrix for all samples, the following pipeline was applied: 1) Trimmed mean of M values (TMM) normalization (*edgeR* package) [85] was applied followed by a 2) Logarithm transformation (apply log(matrix+1)), then 3) Truncation by the quantile 0.99 (all values greater than quantile 0.99 are truncated to this value, all values lower than quantile 0.01 are truncated to this other value) and finally 4) Quantiles normalization (*preprocessCore* package) [86].

### Mechanistic model of cell functionality

The normalized gene expression data was rescaled from the range of variation to 0–1 interval range [max(matrix)=1, min(matrix)=0]. The *Hipathia* method [21], as implemented in the *Hipathia* Bioconductor package [46], was used to estimate signaling circuit activities within the expanded FA pathway from the corresponding normalized gene expression values. The *Hipathia* method uses a Wilcoxon test was used to assess differences in pathway activity between controls and FA samples [21].

### Machine learning

Here, a Multi-Output Random Forest (MORF) regressor that predicts the circuit activity across the whole disease pathway has been implemented using the *scikit-learn* general Machine Learning library [87]. In the learning framework used, the multiple dependent variables that conform the disease environment are modeled in a “all at once” fashion, i.e. each signaling circuit activity in the expanded FA pathway is a target/output variable, whereas each expression value of a KDT gene is an input (Multiple Input Multiple Output). In order to find a “quasi-optimal” set of hyperparameters for our MORF model, we have implemented an optimization strategy on top of *scikit-learn* [87] and *hyperopt* [88]. Since the best hyperparameters to fit the data are problem-dependent [89], the hyperparameter space is explored by means of the TPE [56] method, where each choice of hyperparameters is a “configuration” in the original algorithm. A global R^2^ score averaged across a K-fold cross-validation partition of the data (k=10) is used as objective function. Finally, to evaluate the performance of the model in an unbiased way, the previously found optimal hyperparameters were fixed and a repeated (N=10) k-fold cross-validation is performed.

The same cross-validation can be used to obtain a distribution of the relevance values that can be used to set a threshold beyond which the relevance values obtained by the ML keep their positions in the rank of relevance (have a stable value).

### Enrichment analysis of most relevant genes

Those genes with a relevance confirmed by the cross-validation procedure were considered relevant and were used to perform an enrichment analysis to evaluate their possible impact on the circuits of the FA pathway triggering FA hallmarks. An enrichment analysis was performed by using *enrichR* algorithm using GO Biological Processes as well as Rare Diseases with *AutoRIF* (Automatic Reference into Function) and *GeneRIF* (Gene Reference into Function) from ARCHS^4^ mining of publicly available data tool to predict enrichment in rare diseases terms [90–92]

## List of abbreviations

FA: Fanconi Anemia
GO: Gene Ontology
GP: Gaussian Processes
KDT: Known Drug Targets
KEGG: Kyoto Encyclopedia of Genes and Genomes
ML: Machine Learning
MOR: Multi-Output Regression
MORF: Multi-Output Random Forest
RF: Random Forest
SOR: Single Output Regression
TMM: Trimmed mean of M values
TPE: Tree of Parzen Estimators
TSM: Tree-structured methods

## Declarations

### Availability of data and material

The data used in this study is publicly available in the corresponding repositories cited in the text. The software used is also publicly available in the corresponding web pages, as cited in the text.

## Authors’ contributions

ME has performed the data collection and the analysis, MPC has collaborated in the analysis of the data and the discussion, CL has carried out the machine learning computations and JD has conceived the work and wrote the manuscript.

## Additional Files

### Additional file 1

Excel file (.xls)

Additional Table 1. All gene drug targets studied obtained from DrugBank database version 5.1.2, ranked by their relevance obtained from MORF modelling.

First column: gene name; second column: gene symbol: third column: Entrez ID; fourth column: relevance; fifth column: DrugBank ID of the drugs targeting the gene.

### Additional file 2

Word file (.docx)

Additional Table 2. Genes in the KEGG FA pathway (hsa03460).

First column: gene name; second column: KEGG ID; third column: gene symbol; fourth column: ENSEMBL ID; fifth column: OMIM ID.

### Additional file 3

Tiff file (.tif)

Additional Figure 3. Distribution of circuit activities in the FA KEGG pathway.

Distribution of activities in the seven circuits of the FA KEGG pathway observed in the comparison between healthy and FA bone marrow cells

### Additional file 4

Excel file (.xls)

Additional Table 4. Drugs targeting most relevant genes (relevance>0.005) in Fanconi Anemia extended pathway, obtained from DrugBank database.

First column: DrugBank ID; second column: drug name; third column: drug description; fourth column: drug status; sixth column: drug Indication

### Additional file 5

Excel file (.xls)

Additional Table 5. Enrichment analysis of the most relevant genes.

First column: term detected in the enrichment analysis; second column: overlap; third column: p-value; fourth column: adjusted p-value; fifth column: Z score; sixth column combined score; seventh column genes annotated to the term.

