## Additional file 1 for "Exploring the druggable space around the Fanconi anemia pathway using machine learning and mechanistic models"

**Additional Table 2**. Fanconi Anemia KEGG pathway genes (hsa03460).

| **GENE NAME** | **KEGG ID** | **SYMBOL** | **ENSEMBL ID** | **OMIM** |
| --- | --- | --- | --- | --- |
| Fanconi Anemia complementation group M | hsa:57697 | FANCM | ENSG00000187790 | 609644 |
| Fanconi Anemia complementation group A | hsa:2175 | FANCA | ENSG00000187741 | 227650 |
| Fanconi Anemia complementation group B | hsa:2187 | FANCB | ENSG00000181544 | 300514 |
| Fanconi Anemia complementation group C | hsa:2176 | FANCC | ENSG00000158169 | 227645 |
| Fanconi Anemia complementation group D2 | hsa:2177 | FANCD2 | ENSG00000144554 | 227646 |
| Fanconi Anemia complementation group I | hsa:55215 | FANCI | ENSG00000140525 | 609053 |
| Fanconi Anemia complementation group E | hsa:2178 | FANCE | ENSG00000112039 | 600901 |
| Fanconi Anemia complementation group F | hsa:2188 | FANCF | ENSG00000183161 | 603467 |
| Fanconi Anemia complementation group G | hsa:2189 | FANCG | ENSG00000221829 | 602956 |
| Fanconi Anemia complementation group L | hsa:55120 | FANCL | ENSG00000115392 | 608111 |
| Partner and localizer of BRCA2 | hsa:79728 | PALB2 | ENSG00000083093 | 114480 |
| Fanconi Anemia core complex associated protein 100 | hsa:80233 | FAAP100 | ENSG00000185504 | 611301 |
| BLM RecQ like helicase | hsa:641 | BLM | ENSG00000197299 | 210900 |
| RecQ mediated genome instability 1 | hsa:80010 | RMI1 | ENSG00000178966 | 610404 |
| RecQ mediated genome instability 2 | hsa:116028 | RMI2 | ENSG00000175643 | 612426 |
| DNA topoisomerase III alpha | hsa:7156 | TOP3A | ENSG00000177302 | 601243 |
| DNA topoisomerase III beta 1 | hsa:8940 | TOP3B | ENSG00000100038 | 603582 |
| ATR interacting protein | hsa:84126 | ATRIP | ENSG00000164053 | 606605 |
| ATR serine/threonine kinase | hsa:545 | ATR | ENSG00000175054 | 210600 |
| Replication protein A 30 kDa subunit | hsa:29935 | RPA4 | ENSG00000204086 | 300767 |
| Replication protein A 70 kDa DNA-binding subunit | hsa:6117 | RPA1 | ENSG00000132383 | 179835 |
| Replication protein A 32 kDa DNA-binding subunit | hsa:6118 | RPA2 | ENSG00000117748 | 179836 |
| Replication protein A 14 kDa subunit | hsa:6119 | RPA3 | ENSG00000106399 | 179837 |
| Telomere length regulation protein TEL2 homolog | hsa:9894 | TELO2 | ENSG00000100726 | 611140 |
| Breast cancer type 2 susceptibility protein | hsa:675 | BRCA2 | ENSG00000139618 | 114480 |
| DNA mismatch repair protein Mlh1 | hsa:4292 | MLH1 | ENSG00000076242 | 120436 |
| Fanconi anemia group J protein | hsa:83990 | BRIP1 | ENSG00000136492 | 114480 |
| Breast cancer type 1 susceptibility protein | hsa:672 | BRCA1 | ENSG00000012048 | 113705 |
| DNA repair protein RAD51 homolog 3 | hsa:5889 | RAD51C | ENSG00000108384 | 602774 |
| Fanconi-associated nuclease 1 | hsa:22909 | FAN1 | ENSG00000198690 | 613534 |
| Ubiquitin carboxyl-terminal hydrolase 1 | hsa:7398 | USP1 | ENSG00000162607 | 603478 |
| WD repeat-containing protein 48 | hsa:57599 | WDR48 | ENSG00000114742 | 612167 |
| Ubiquitin-conjugating enzyme E2 T | hsa:29089 | UBE2T | ENSG00000077152 | 610538 |
| REV1, DNA directed polymerase | hsa:51455 | REV1 | ENSG00000135945 | 606134 |
| CENPS-CORT readthrough | hsa:100526739 | CENPS-CORT | ENSG00000251503 | - |
| Centromere protein X | hsa:201254 | CENPX | ENSG00000169689 | 615128 |
| Centromere protein S | hsa:378708 | CENPS | ENSG00000175279 | 609130 |
| DNA polymerase eta | hsa:5429 | POLH | ENSG00000170734 | 278750 |
| DNA polymerase kappa | hsa:51426 | POLK | ENSG00000122008 | 605650 |
| REV3 like, DNA directed polymerase zeta catalytic subunit | hsa:5980 | REV3L | ENSG00000009413 | 602776 |
| DNA polymerase iota | hsa:11201 | POLI | ENSG00000101751 | 605252 |
| PMS1 homolog 2, mismatch repair system component | hsa:5395 | PMS2 | ENSG00000122512 | 276300 |
| MUS81 structure-specific endonuclease subunit | hsa:80198 | MUS81 | ENSG00000172732 | 606591 |
| Essential Meiotic Structure-Specific Endonuclease Subunit 1 | hsa:146956 | EME1 | ENSG00000154920 | 610885 |
| Essential Meiotic Structure-Specific Endonuclease Subunit 2 | hsa:197342 | EME2 | ENSG00000197774 | 610886 |
| ERCC excision repair 4, endonuclease catalytic subunit | hsa:2072 | ERCC4 | ENSG00000175595 | 133520 |
| ERCC excision repair 1, endonuclease non-catalytic subunit | hsa:2067 | ERCC1 | ENSG00000012061 | 126380 |
| SLX1 homolog A, structure-specific endonuclease subunit | hsa:548593 | SLX1A | ENSG00000132207 | 615822 |
| SLX1 homolog A, structure-specific endonuclease subunit | hsa:79008 | SLX1B | ENSG00000181625 | 615823 |
| SLX4 structure-specific endonuclease subunit | hsa:84464 | SLX4 | ENSG00000188827 | 613278 |
| RAD51 recombinase | hsa:5888 | RAD51 | ENSG00000051180 | 114480 |
| FA core complex associated protein 24 | hsa:91442 | FAAP24 | ENSG00000131944 | 610884 |
| Hes family bHLH transcription factor 1 | hsa:3280 | HES1 | ENSG00000114315 | 139605 |
| DNA polymerase nu | hsa:353497 | POLN | ENSG00000130997 | 610887 |
