## Supplementary figures and images for "Exploring the druggable space around the Fanconi anemia pathway using machine learning and mechanistic models"

### Additional file 2

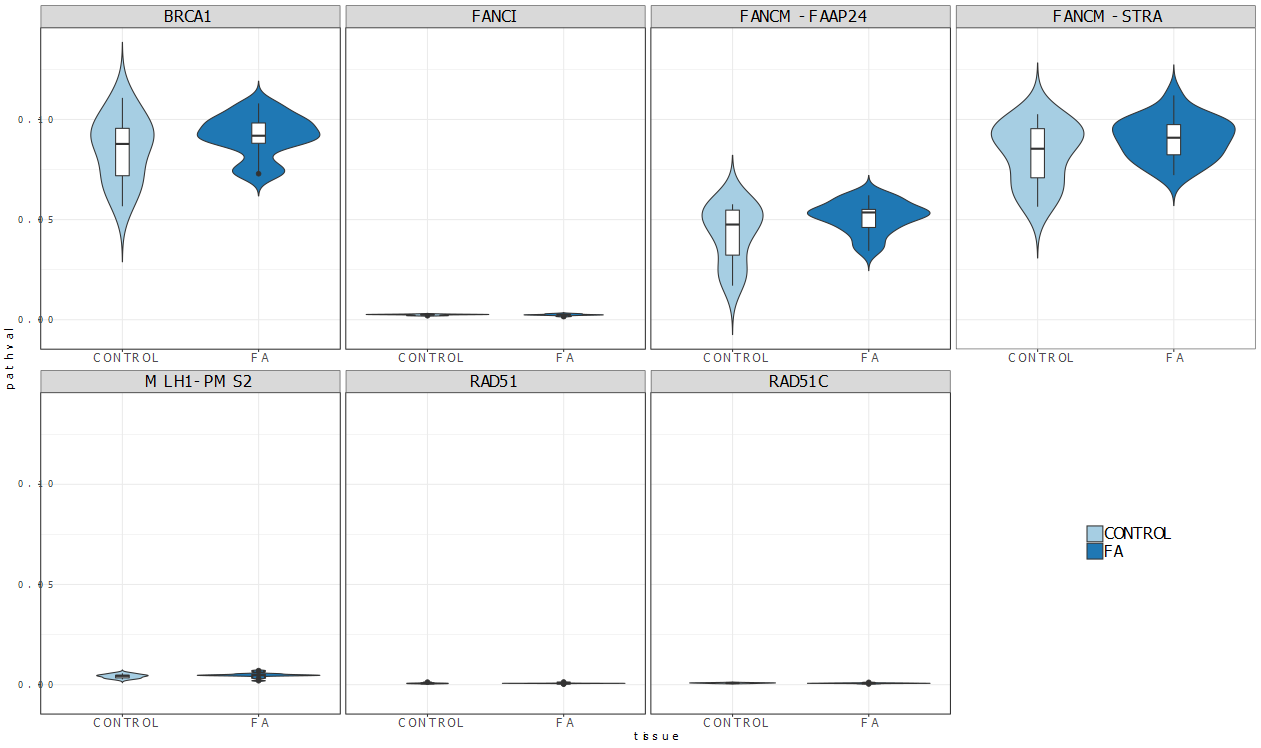
